# Immunomodulatory actions of a kynurenine-derived endogenous electrophile

**DOI:** 10.1101/2022.02.08.479451

**Authors:** Mara Carreño, Maria F. Pires, Steven R. Woodcock, Tomasz Brzoska, Samit Ghosh, Sonia R. Salvatore, Matthew Dunn, Nora Connors, Shuai Yuan, Adam C. Straub, Stacy G. Wendell, Gregory J. Kato, Bruce A. Freeman, Solomon Ofori-Acquah, Prithu Sundd, Francisco J. Schopfer, Dario A. Vitturi

## Abstract

The inflammatory upregulation of kynurenine metabolism induces immunomodulatory responses via incompletely understood mechanisms. We report that increases in cellular and systemic kynurenine levels yield the electrophilic derivative kynurenine-carboxyketoalkene (Kyn-CKA), as evidenced by the accumulation of thiol-conjugates and saturated metabolites. Under physiological conditions, Kyn-CKA induces Nrf2-regulated genes and inhibits NF-κB and NLRP3-dependent pro-inflammatory signaling. Sickle Cell Disease (SCD) is a hereditary hemolytic condition characterized by basal inflammation and recurrent vaso-occlusive crises. Both a transgenic SCD murine model and SCD patients exhibit increased kynurenine synthesis and elevated Kyn-CKA metabolite levels. Plasma hemin and kynurenine concentrations are positively correlated, indicating that Kyn-CKA synthesis in SCD is upregulated during pathogenic vascular stress. Remarkably, exogenous administration of Kyn-CKA abrogated pulmonary microvasculature occlusion in SCD mice, an important factor in the development of lung injury. These findings demonstrate that the upregulation of kynurenine synthesis and its metabolism to Kyn-CKA is an adaptive response that attenuates inflammation and protects tissues.

**One-Sentence Summary:** Kyn-CKA is a kynurenine-derived signaling mediator that transduces its immunomodulatory protective actions and attenuates vaso-occlusion in sickle cell disease.

## Introduction

Acute and chronic inflammation induces the rate-limiting enzymes indoleamine dioxygenase-1 (IDO1) and tryptophan dioxygenase-2 (TDO2) that in turn increase kynurenine and kynurenine/tryptophan ratios (*1*). Increased kynurenine synthesis via TDO2 in hepatocytes is protective against primary endotoxemia, while IDO1 induction in myeloid leukocytes attenuates inflammatory responses to endotoxin, promotes the generation of tolerogenic dendritic cells (DC) and facilitates T_reg_ polarization (*2–5*). Consistent with these immunomodulatory effects, dysregulated kynurenine metabolism occurs in inflammatory conditions such as cardiovascular disease, cancer, renal injury, transplantation and is often a strong predictor of outcome (*6–9*). However, despite its pathophysiological relevance, the mechanisms responsible for the immunoregulatory effects of kynurenine and its metabolites are poorly defined (*10*).

Biological electrophiles contain carbon-carbon double bonds conjugated to electron-withdrawing moieties that generate electron-deficient carbons that are susceptible to nucleophilic attack by thiols or, less predominantly, amines (*11*). Covalent modification of critical cysteine thiols in transcription factors, signaling proteins, and metabolic enzymes have profound effects on cellular function (*11*). In particular, reactive electrophiles elicit cytoprotective and immunomodulatory effects both preclinically and clinically by upregulating Nrf2-dependent antioxidant enzyme translation and inhibiting NF-κB-dependent gene expression and inflammatory cell metabolic reprogramming (*11, 12*). *In vivo,* kynurenine deamination generates an α,β-unsaturated carbonyl-containing product, Kynurenine-Carboxy-Keto-Alkene (Kyn-CKA), which participates in the generation of UV-filters in the eye lens (*13*). Critically, the potential for Kyn-CKA to contribute to the immunomodulatory actions of kynurenine metabolites has not been explored.

Herein, we show that kynurenine synthesis in hepatocytes and macrophages generates Kyn-CKA. Novel Kyn-CKA-specific metabolites detected in healthy humans and mice affirms that Kyn-CKA formation is biologically significant, with Kyn-CKA generation upregulated by increases in systemic kynurenine concentrations. Kyn-CKA activates Nrf2-dependent gene expression and inhibits NF-κB and NLRP3-dependent pro-inflammatory responses in both parenchymal and immune cells. These responses are recapitulated across several tissues *in vivo*. In addition to upregulating Nrf2-dependent signaling under basal non-activated conditions, Kyn-CKA attenuates LPS-induced pro-inflammatory gene expression and cytokine synthesis. Finally, Kyn-CKA increases IL-10 levels and attenuates the expression of renal injury markers in LPS-treated mice, suggesting potent anti-inflammatory and cytoprotective actions *in vivo*.

The anti-inflammatory effects of Kyn-CKA suggested the possibility that its formation might constitute a missing link in kynurenine signaling and an adaptive immunomodulatory response having the potential to modulate disease progression. To test this hypothesis, we focused on the hemolytic inflammatory conditions intrinsic to Sickle Cell Disease (SCD), wherein the activation of endothelial, platelet and leukocyte Toll-like Receptor-4 (TLR4) by hemoglobin-derived products promotes the formation of multicellular aggregates which, in combination with increased sickle erythrocyte rigidity and adhesiveness, cause vaso-occlusion and ischemic injury. This phenomenon, not only causes severe pain crises but also, leads to the development of lifethreatening acute and chronic complications in SCD patients (*14, 15*). Analysis of plasma samples obtained from a transgenic humanized SCD mouse model and from SCD patients showed elevated kynurenine levels which correlated positively with pro-inflammatory plasma hemin concentrations. This increased kynurenine synthesis was further correlated with greater Kyn-CKA metabolite concentrations in the plasma and urine of SCD mice and SCD patients. Quantitative fluorescence intravital microscopy was used to monitor vaso-occlusive occurrence in the lung microvasculature of SCD mice and to test the role of Kyn-CKA as a modulator of tissue dysfunction (*16, 17*). Consistent with a role Kyn-CKA as an endogenous antiinflammatory mediator in SCD, Kyn-CKA significantly decreased the average size of multicellular aggregates and inhibited vaso-occlusion of the pulmonary microvasculature.

Collectively, our results reveal that Kyn-CKA is present under basal conditions and that its formation is adaptively elevated within the context of inflammatory kynurenine synthesis upregulation. Kyn-CKA is an endogenous electrophilic signaling mediator capable of attenuating pro-inflammatory signaling cascades, modulating complex pathological responses, and potentially, impacting the course of human disease.

## Results

### Kynurenine yields Kyn-CKA and promotes Nrf2-dependent gene expression

Kynurenine is a main product of tryptophan (Trp) metabolism and, upon deamination, is activated into Kyn-CKA, an α,β-unsaturated carbonyl-containing derivative with electrophilic reactivity (Fig 1A). Treatment of the normal mouse hepatocyte cell line AML-12 with Kyn-CKA dose-dependently upregulated electrophile-sensitive Nrf2-dependent gene expression both at the transcript and protein levels, suggesting that Kyn-CKA is a chemically and biologically active product (Fig 1B,C). Consistent with its electrophilic reactivity, Kyn-CKA rapidly targeted intracellular thiols, as evidenced by the formation of glutathione (GSH) conjugates (GSH-Kyn-CKA) that were excreted and detected in the extracellular milieu (Fig 1D). To define the contribution of intracellular enzymatic reactions to the endogenous formation of Kyn-CKA, AML-12 were treated with Trp and the levels of intermediate metabolites N-formyl-kynurenine and kynurenine were assessed. Trp caused a significant increase in N-formyl-kynurenine and kynurenine synthesis that led to the formation of GSH-Kyn-CKA adducts (Fig 1E-G). Similarly, treatment with N-formyl-kynurenine (NF-Kyn) or L-kynurenine (L-Kyn) resulted in dose-dependent GSH-Kyn-CKA formation (Fig 1H). The formation of GSH conjugates is the first step in the mercapturic acid pathway, a conserved mechanism for cellular electrophile inactivation that is followed by export through multidrug resistance proteins (MRP). Consistent with this pathway being engaged in Kyn-CKA metabolism, treatment with the pan-MRP inhibitor probenecid induced a significant increase in intracellular GSH-Kyn-CKA levels (Fig 1I). Finally, and parallel to the endogenous formation of GSH-Kyn-CKA adducts, treatment of AML-12 cells with Trp significantly induced Keap1-regulated Nrf2-dependent gene expression (Fig 1J). This underscores that intracellular Kyn-CKA formation can evade both high cellular GSH concentrations and trapping by the mercapturic acid pathways to induce Keap1 thiol alkylation and Nrf2-regulated signaling. Consistent with its high electrophilic reactivity, non-thiol adducted Kyn-CKA was not detectable in both control and Trp-treated cells (not shown).

**Fig. 1.**
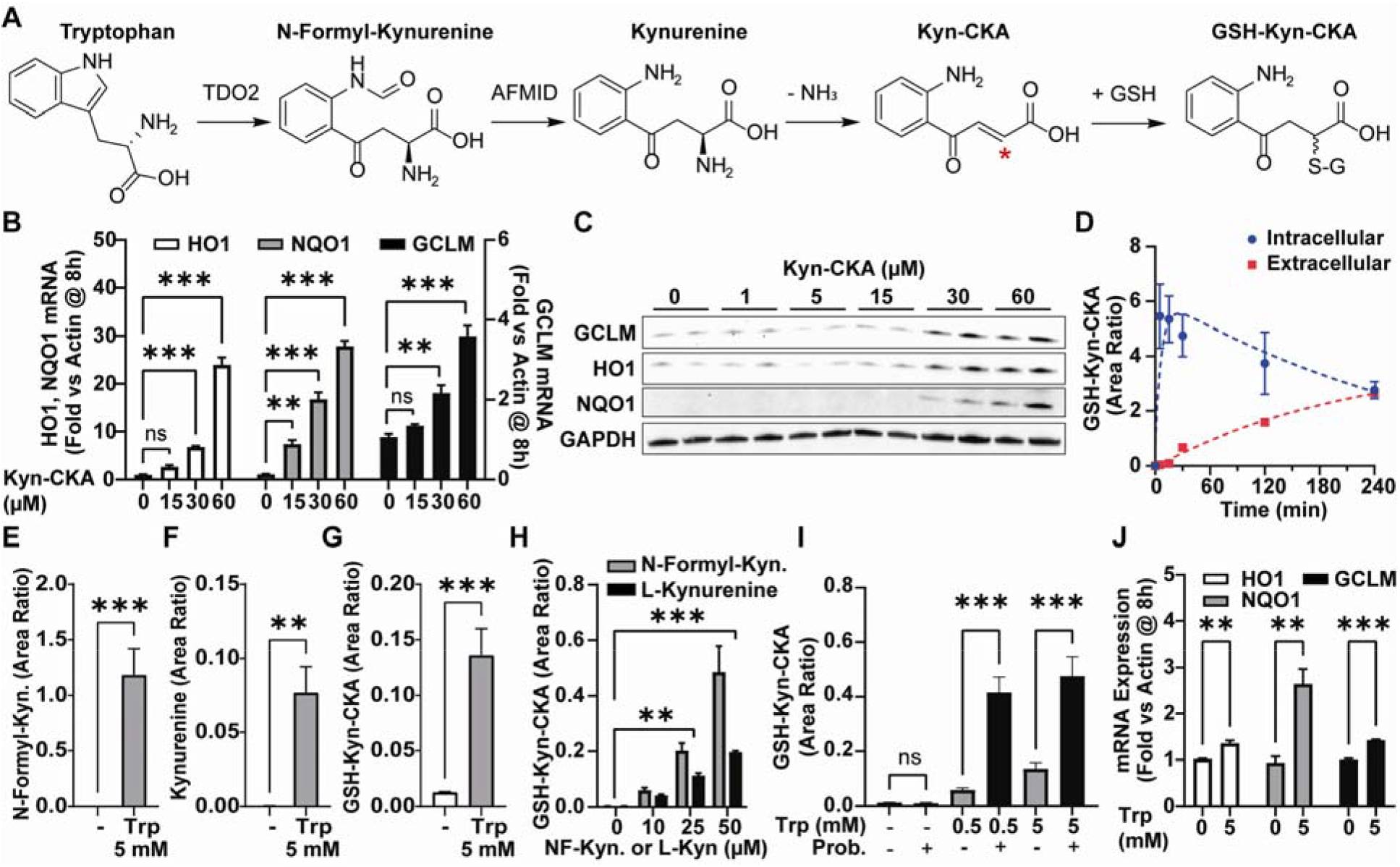
Bioactive Kyn-CKA is formed following kynurenine synthesis in AML-12 cells. (**A**) Proposed route for Kyn-CKA formation following from Trp metabolism. The asterisk denotes the electrophilic carbon. (B) Nrf2-dependent gene induction by Kyn-CKA (n=3). (C) Nrf2-regulated protein expression 24 h post Kyn-CKA. (D) GSH-Kyn-CKA formation following Kyn-CKA treatment (60 μM, n=2 per time point). Data is expressed as the ratio between GSH-Kyn-CKA and Trp-d_3_ areas. (E-G) Kynurenine metabolites 8 h post Trp (5 mM, n=3). (H) GSH-Kyn-CKA formation 8 h post NF-Kyn or L-Kyn (n=2 per dose). (I) Effect of probenecid (1 mM) on GSH-Kyn-CKA after Trp for 8 h (n=3). (J) Nrf2-dependent gene expression 8 h post Trp (n=3). ** p < 0.01, ***p < 0.0001 by one- (B) or two-way ANOVA and Tuckey’s test (I), or by t-test (E-H,J).

### In vivo generation of Kyn-CKA

*In vivo*, intracellular electrophile-GSH conjugates are exported and metabolized to yield cysteine and mercapturic acid conjugates that are excreted in urine. To evaluate Kyn-CKA-specific metabolites *in vivo,* urine was collected from C57Bl/6J mice either in the absence or 8 h following Kyn-CKA supplementation (20, 50 mg/kg ip). Consistent with its endogenous nature, LC-MS/MS analyses demonstrated the presence of urinary Cys-Kyn-CKA adducts in untreated mice which was confirmed by co-elution with a synthetic ^13^C_3-_Cys-Kyn-CKA standard (Fig 2A). Exogenous Kyn-CKA administration gave higher levels of urinary Cys-Kyn-CKA adducts and led to the excretion of N-acetyl-cysteine Kyn-CKA conjugates (NAC-Kyn-CKA, Fig 2A,B and Suppl Fig 1). In addition, liver tissue analysis revealed endogenous GSH-Kyn-CKA adducts in untreated mice, with GSH-Kyn-CKA significantly increased by Kyn-CKA treatment (Fig 2C,D and Suppl Fig 1). These results were recapitulated in C57Bl/6J treated with L-Kyn, suggesting that systemic increases in this amino acid metabolite can promote Kyn-CKA formation *in vivo* (Fig 2E,F). Untargeted analysis of plasma samples from Kyn-CKA-treated mice revealed unadducted Kyn-CKA together with several upregulated LC-MS/MS features, including a reduced carboxyketoalkane derivative termed Red-Kyn-CKA (Fig 2G, Suppl Fig 2). Red-Kyn-CKA formation from Kyn-CKA was validated using a rat liver S9 fraction plus NADPH as a source of reducing equivalents (Suppl Fig 2) and confirmed by comparing the MS^2^ fragmentation pattern with a synthetic standard (Fig 2H). Unlike urinary and hepatic conjugates, neither free Kyn-CKA nor Red-Kyn-CKA were detected in untreated mouse plasma (not shown). Transcript expression analysis from Kyn-CKA treated mice revealed induction of Nrf2-dependent signaling in kidneys, heart, and liver (Fig 2I-K) and confirmed at the protein level in liver and kidneys (Fig 2L,M). No changes were observed in lung and brain (not shown). Nrf2-dependent gene expression was also upregulated in the liver of L-Kyn treated mice, consistent with Kyn-CKA formation in this tissue (Fig 2N) and reinforced by both increased levels of hepatic GSH-Kyn-CKA and urinary Cys-Kyn-CKA conjugates (Fig 2E,F).

**Fig. 2.**
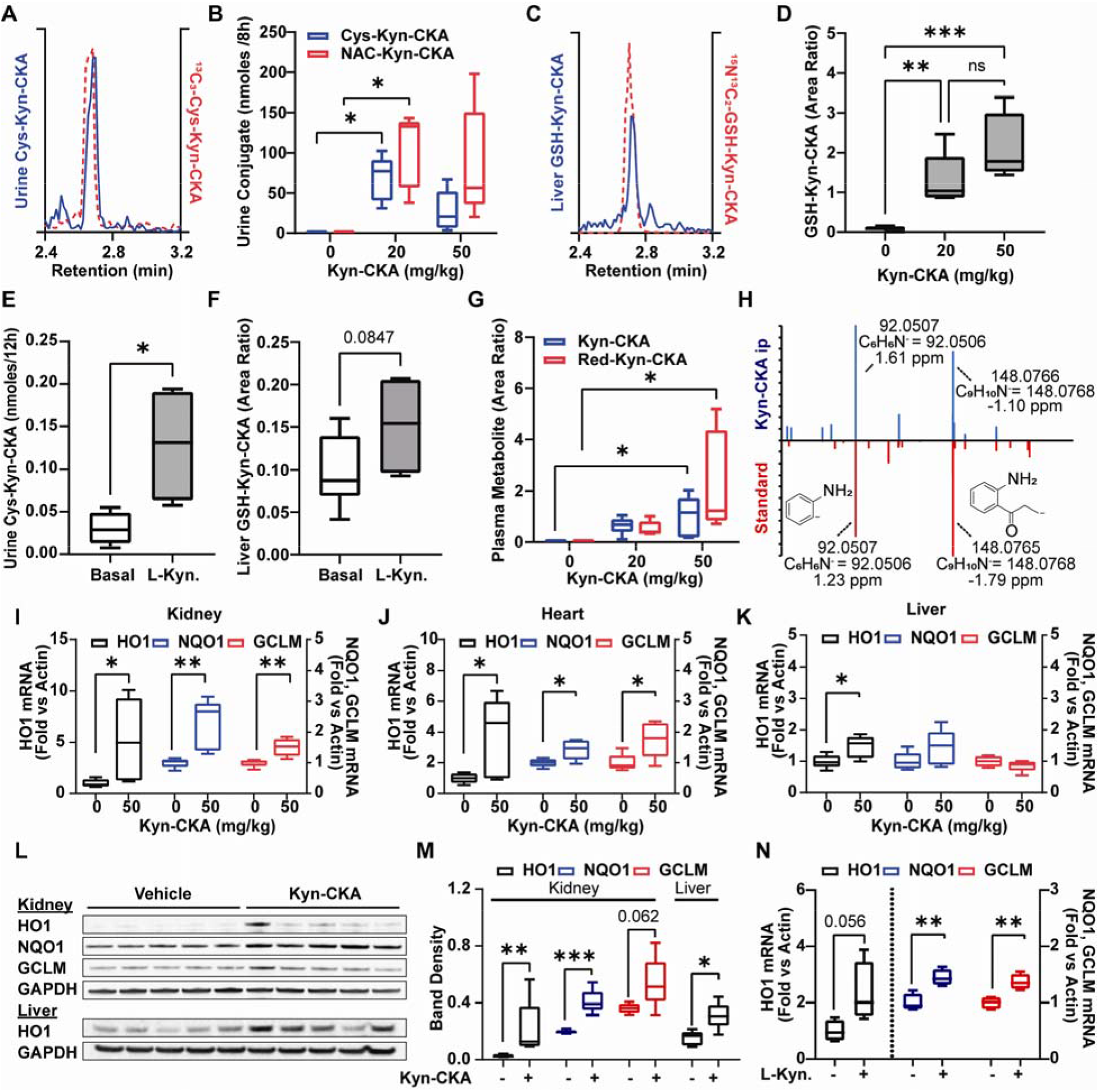
Kyn-CKA is formed endogenously and induces Nrf2-dependent genes *in vivo.* (**A**) Representative LC-MS/MS trace showing co-elution between endogenous urinary Cys-Kyn-CKA and an isotopically labeled synthetic standard. (B) Urine Cys- and NAC-Kyn-CKA 8 h post Kyn-CKA (ip) in C57Bl6/J mice (n=5 per dose). (C) LC-MS/MS trace of endogenous GSH-Kyn-CKA in untreated mouse liver. (D) Hepatic GSH-Kyn-CKA in mice 8 h following Kyn-CKA (basal n=9, n=5 per dose). (E) Urinary Cys-Kyn-CKA in mice receiving L-Kyn (100 mg/kg ip, 2 d, n=4). (F) Hepatic GSH-Kyn-CKA L-Kyn treated mice (100 mg/kg ip, 2 d, basal n=9, L-Kyn n=4). (G) Plasma Kyn-CKA metabolites 8 h post-dosing. (H) Red-Kyn-CKA MS^2^ spectra for Kyn-CKA treated plasma (20 mg/kg, above) and synthetic Red-Kyn-CKA (below). (I-K) Nrf2-dependent gene expression in mice 8 h post Kyn-CKA (50 mg/kg, n=5 per dose). (L, M) Nrf2-dependent protein levels in mice 24 h post Kyn-CKA (20 mg/kg ip, 2 doses, 8 h apart). (N) Nrf2-dependent gene expression in mice 12 h post second L-Kyn dose (50 mg/kg ip, 2 doses, 24 h apart, n=4). * p < 0.05, ** p < 0.01, *** p < 0.0001 by one-way ANOVA and Tukey’s test (B,D,G) or t-test (E,F,I-N). Kidney HO-1 in (M) was analyzed by Mann-Whitney test.

### Kyn-CKA inhibits NF-κB-dependent signaling and NLRP3 inflammasome engagement in macrophages

While TDO2 catalyzes kynurenine synthesis in the liver, this reaction is mediated by IDO1 in myeloid cells. Akin to the results obtained with AML12 hepatocytes, Trp treatment of J774a. 1 macrophages increased the formation of N-formyl-kynurenine and kynurenine as well as GSH-Kyn-CKA (Fig 3A,C). In addition to inducing Nrf2-dependent gene expression (Suppl. Fig 3A), Kyn-CKA inhibited the LPS-induced expression of pro-inflammatory cytokines and enzymes that generate nitrogen oxides in both J774a. 1 (Fig 3D-H) and bone marrow-derived macrophages (BMDM, Suppl. Fig 3B,C). NF-κB activation is essential in priming the NLRP3 inflammasome via increased synthesis of NLRP3 subunits and pro-IL-1β. Activation of the primed inflammasome promotes proteolytic pro-caspase-1 activation followed by mature IL-1β generation (*12*). Kyn-CKA treatment of LPS/ATP activated J774a.1 dose-dependently inhibited NLRP3 and pro-IL1β expression and attenuated pro-caspase-1 levels to inhibit processing and secretion of mature IL-1β (Fig 3I), with similar results obtained using BMDM (Suppl Fig 3D). Beyond an effect on myeloid cells, Kyn-CKA inhibited LPS-induced VCAM-1 expression in human pulmonary microvascular endothelial cells (HPMVEC), supporting that Kyn-CKA can invoke systemic anti-inflammatory responses across different cell types and tissues (Fig 3J).

**Fig. 3.**
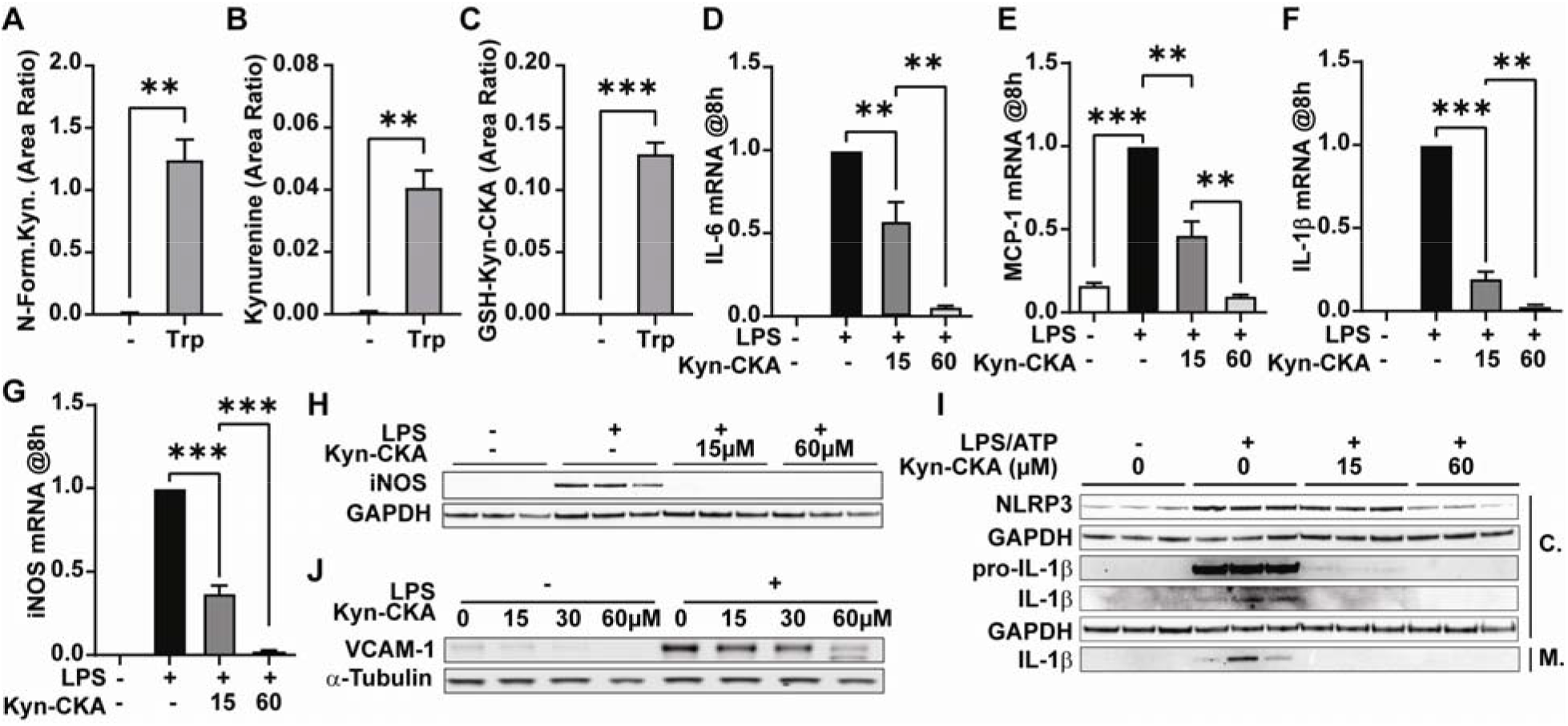
Kyn-CKA is formed and attenuates NF-kB and NLRP3 signaling in macrophages. (**A-C**) Kynurenine metabolites in J774a.1 4 h post Trp (5 mM, n=3). (D-G) NF-κB-dependent gene expression 8 h post LPS (100 ng/mL) and Kyn-CKA (15 or 60 μM, n=3). (H) iNOS expression 8 h post LPS (100 ng/mL) and Kyn-CKA. (I) Inhibition of NLRP3 inflammasome activation by LPS/ATP (100 ng/mL, ATP 2 mM) by Kyn-CKA at 8 h. C: Cellular fraction, M: Media. (J) VCAM-1 expression in HPMVEC 16 h post LPS (100 ng/mL) and Kyn-CKA. ** p < 0.01, *** p < 0.0001 by t-test (A-C) or one-way ANOVA and Tukey’s test (D-G).

To determine whether the anti-inflammatory effects of Kyn-CKA *in vitro* translate to *in vivo* inflammatory scenarios, C57Bl6/J mice were treated with a single dose of Kyn-CKA (50 mg/kg ip) followed by LPS challenge 30 min later (10 mg/kg ip). Tissue NF-κB-dependent gene expression revealed anti-inflammatory responses to Kyn-CKA in kidneys (Fig 4A-D), heart (Fig 4E-H), and lung (Fig 4I). Kyn-CKA pretreatment also decreased the expression of the renal injury marker Kim-1, affirming attenuation of LPS-induced acute kidney injury (Fig 4J). Consistent with the induction of systemic anti-inflammatory responses, Kyn-CKA significantly inhibited plasma levels of several pro-inflammatory cytokines (Fig 4K-O) while increasing the anti-inflammatory mediator IL-10 (Fig 4P).

**Fig. 4.**
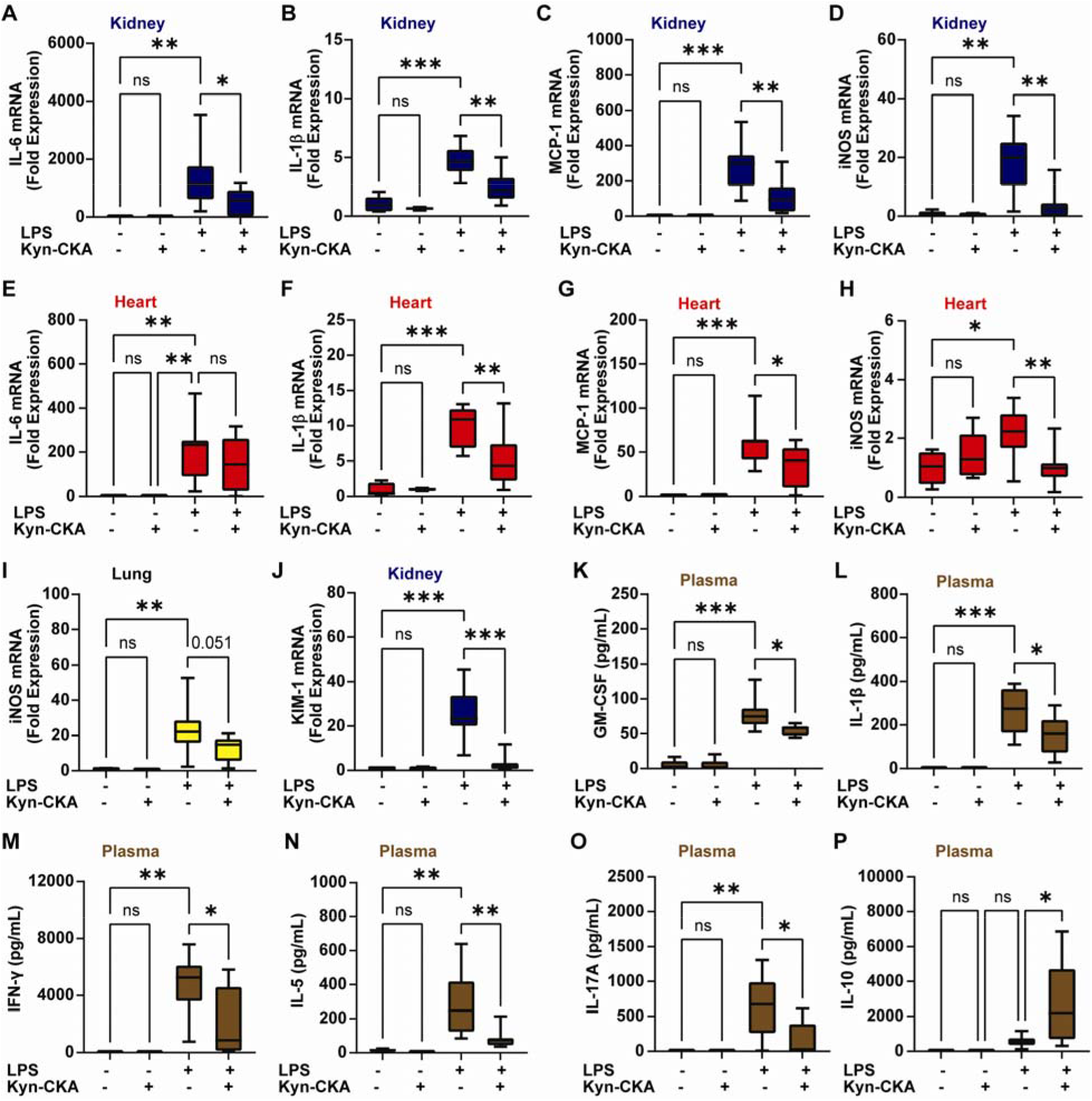
Kyn-CKA promotes anti-inflammatory actions *in vivo.* (**A-I**) Kyn-CKA (50 mg/kg ip, 30 min before challenge) inhibits NF-κB target expression 8 h post LPS (10 mg/kg ip). (J) Kyn-CKA inhibits 8 h Kim-1 following LPS. (K-P) Plasma cytokine levels modulated by Kyn-CKA 8 h post-LPS. Non-LPS treated (n=5), LPS-treated (n=8-9). *p < 0.05, **p < 0.01, ***p < 0.001 by one-way ANOVA and Tukey’s test.

### Kyn-CKA levels are elevated in SCD and Kyn-CKA administration attenuates inflammatory vaso-occlusion

To define the potential contributions of Kyn-CKA formation under clinically relevant pathological conditions, the concentrations of Kyn-CKA metabolites were determined in a humanized mouse model of SCD. Compared to non-sickle HbAA controls, homozygous HbSS mice exhibited higher hepatic TDO2 expression and elevated plasma kynurenine levels (Fig 5A-C). Interestingly, a strong positive correlation was observed between kynurenine and hemin concentrations in HbSS mouse plasma, indicating an association between pathogenic hemin accumulation and kynurenine synthesis (Fig 5D). In this regard, while hemin failed to induce TDO2 in AML12 hepatocytes (not shown), it upregulated IDO1 expression in J774a.1 (Fig 5E). Consistent with increased Kyn-CKA formation secondary to kynurenine synthesis, urinary Cys-Kyn-CKA levels were significantly greater in HbSS versus HbAA mice (Fig 5F). This was recapitulated clinically in a cohort of SCD patients that, in addition to greater plasma kynurenine concentrations, also exhibited increased levels of both urinary Cys-Kyn-CKA and plasma Red-Kyn-CKA (Fig 5G-I). No differences in circulating tryptophan levels were observed between SCD and non-SCD subjects in either mouse or human cohorts, and kynurenine to tryptophan ratios were consistent with kynurenine pathway upregulation in SCD (Suppl Fig 4).

**Fig. 5.**
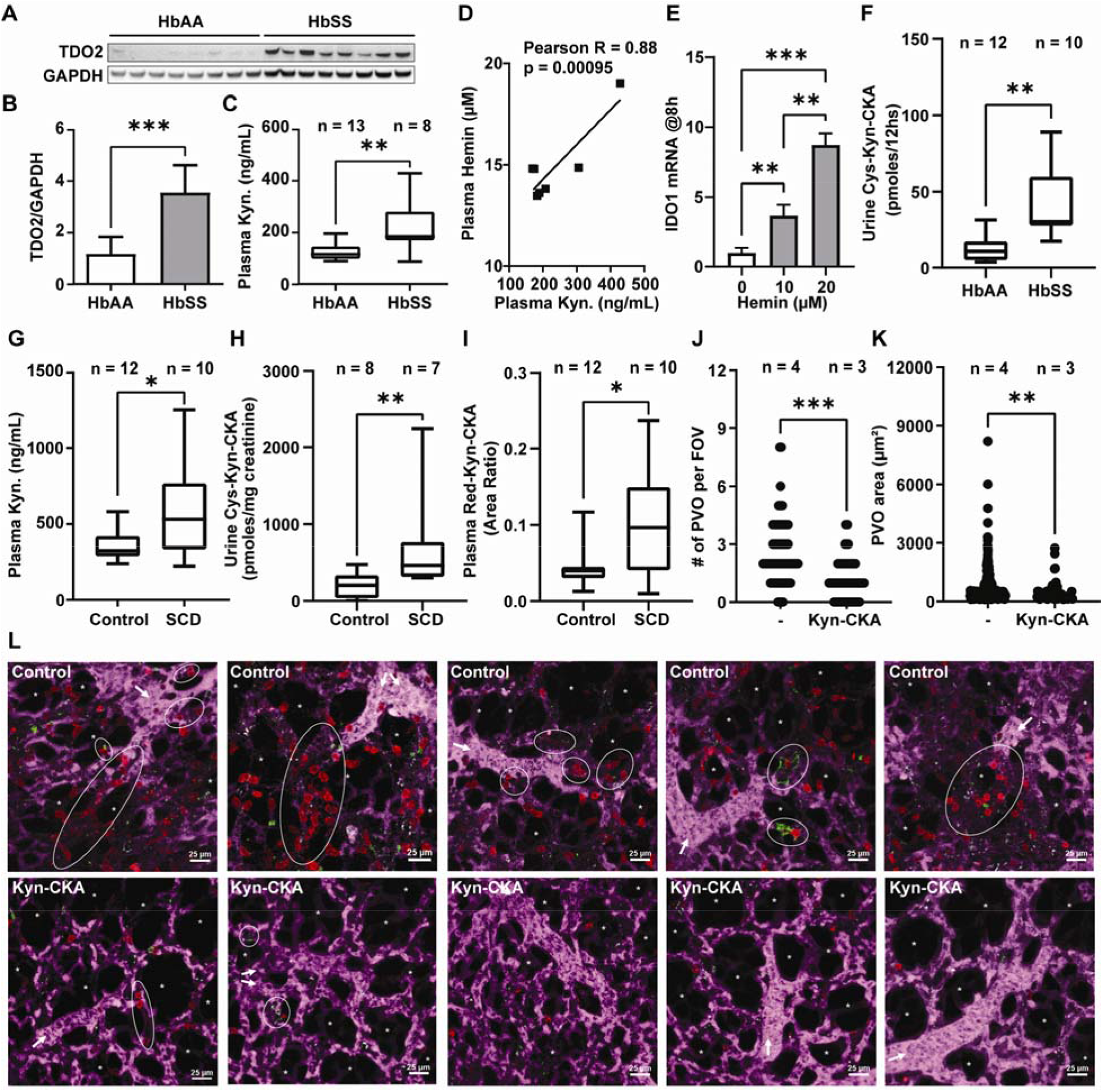
Kyn-CKA synthesis is elevated in SCD and attenuates pulmonary microvascular vaso-occlusion. (**A-B**) TDO2 immunoblot and quantification (n=8 per genotype). (C) Plasma kynurenine in HbAA vs HbSS mice. (D) Correlation between HbSS plasma kynurenine and hemin (n = 7). (E) IDO1 induction by hemin in J774a.1 (n=3 per condition). (F). Cys-Kyn-CKA in 12 h HbAA and HbSS urine. (G-I) Plasma kynurenine, urine Cys-Kyn-CKA and plasma Red-Kyn-CKA in healthy and SCD volunteers. (J-K) Number of pulmonary vaso-occlusions per field of view and pulmonary vaso-occlusion area in HbSS mice receiving PBS or Kyn-CKA (10 mg/kg iv) 30 min prior to LPS (0.1 μg/kg iv). Control n = 4 (89 FOVs), Kyn-CKA n = 3 (60 FOVs). (L) Representative fields of view for HbSS mice treated as in K. Asterisks denote alveolar spaces, arrows indicate blood flow direction and dotted ellipses show pulmonary vasoocclusions. * p < 0.05, ** p < 0.01, *** p < 0.0001 by t-test (A-C, G, I), Mann Whitney test (F, H, J, K) or one-way ANOVA and Tukey’s test (E).

To test whether the anti-inflammatory actions of Kyn-CKA can modulate SCD-specific pathology, inflammatory vaso-occlusion was monitored in real-time by intravital microscopy of the pulmonary microvasculature (*16, 17*). Intravenous injection of a nanogram dose of LPS (0.1 μg/kg) in HbSS mice promoted formation of multicellular aggregates comprised of platelets and neutrophils that obstructed pulmonary arterioles and led to the interruption of blood flow in the lung. Pretreatment with Kyn-CKA (10 mg/kg, iv) significantly inhibited both pulmonary vasoocclusion and the size of individual multicellular aggregates (Fig 5J-L and Suppl Media). These results reinforce that Kyn-CKA formation secondary to kynurenine synthesis upregulation in SCD is an adaptive anti-inflammatory response that inhibits inflammatory events leading to vaso-occlusion and protects organs against the cumulative effects of recurrent metabolic and physical stresses.

## Discussion

The upregulation of the rate-limiting enzymes in kynurenine metabolism is linked with diverse anti-inflammatory and immunomodulatory effects, but the precise mechanisms behind these immunosuppressive actions are incompletely understood (*10*). Proposed effects include TDO2- or IDO1-mediated decreases in tryptophan, alterations in NAD^+^ availability, activity-independent signaling actions of IDO1, and the generation of bioactive kynurenine metabolites (*2, 10, 18*). In particular, the activation of the aryl hydrocarbon receptor (AhR) and the G-protein coupled receptor 35 (GPR35) by kynurenine and kynurenic acid have been proposed, but whether their physiological concentrations are sufficient to engage these pathways *in vivo* is controversial (*19, 20*). Herein, we demonstrate that kynurenine yields a bioactive electrophile, Kyn-CKA, *in vitro* and *in vivo.* Moreover, the anti-inflammatory signaling actions of Kyn-CKA suggest that this novel metabolite could account for a significant fraction of the immunomodulatory effects ascribed to the kynurenine pathway.

A facile reaction with abundant nucleophilic thiols and amines, combined with substantial metabolic transformation, challenges the *in vivo* detection of endogenous electrophiles in their “free” non-adducted form, as is the case for Kyn-CKA. Strategies for electrophile discovery involve the use of highly specific LC-MS/MS and LC-HRMS detection of proximal metabolites such as non-electrophilic reduction products and excreted urinary conjugates. This approach allowed the detection of several specific Kyn-CKA metabolites *in vivo:* the non-electrophilic Red-Kyn-CKA in plasma, an intracellular GSH-Kyn-CKA conjugate, and the corresponding cysteine- and N-acetyl-cysteine adducts in urine. Unequivocal identification of these metabolites is supported by co-elution with isotopically labeled standards for thiol-Kyn-CKA adducts, confirmation of molecular composition, and fragmentation analyses at the <2 ppm level using LC-HRMS. Importantly, the observation that Red-Kyn-CKA, Cys-Kyn-CKA, and GSH-Kyn-CKA are present in plasma, urine, and liver extracts from untreated mice and healthy humans demonstrate that Kyn-CKA is an endogenous mediator produced *in vivo* under normal physiological conditions.

Electrophilic molecules such as Kyn-CKA covalently modify hyperreactive and functionally significant cysteine thiols in transcription factors, signaling proteins and enzymes to induce changes in protein and cellular function. Nucleophilic targets susceptible to electrophilic addition reactions include critical cysteines of TLR/NF-κB signaling, NLRP3 inflammasome components, and the Nrf2 inhibitor protein Keap1 (*11, 12*). Herein, we show both *in vitro* and *in vivo* that Kyn-CKA activates the expression of Nrf2-dependent proteins and attenuates the production of pro-inflammatory cytokines by inhibiting TLR4 and NLRP3 engagement. In the kidney, Kyn-CKA induced Nrf2-regulated gene expression, attenuated pro-inflammatory genes, and decreased the expression of the renal injury marker Kim-1 upon LPS challenge. These observations are consistent with the role of the kidney in a) kynurenine reabsorption and metabolism, b) Kyn-CKA excretion as urinary Cys-Kyn-CKA conjugates, and c) the protective effect of Nrf2 inducers against diverse renal injuries (*21, 22*). Having demonstrated the endogenous nature and robust signaling actions of Kyn-CKA, it is also shown that Kyn-CKA synthesis is upregulated under clinically relevant inflammatory conditions and impacts complex pathological responses. SCD was selected because of its chronic inflammatory nature, reports of elevated plasma kynurenine levels in patients, and a precedent for reactive electrophiles being able to modulate disease outcome (*15, 23, 24*). In line with published results (*23*), plasma kynurenine levels were elevated in both a murine SCD model and SCD patients, which was in turn associated with increased urinary Cys-Kyn-CKA and plasma Red-Kyn-CKA. Moreover, a strong positive correlation was observed between hemin and kynurenine levels in plasma, indicating that pathologic events in SCD promote kynurenine pathway upregulation possibly through a combination of direct and indirect effects of hemin on IDO1 and TDO2 expression (*2, 25*). The heightened inflammatory state of SCD results from upregulated TLR4 expression, release of pro-inflammatory hemoglobin-derived products during intravascular hemolysis, and increased gut endotoxin leakage (*26–29*). In this context, TLR4 activation by hemin and LPS in endothelial and immune cells and NLRP3 inflammasome engagement in platelets are critical to the onset of vaso-occlusive crises (*17, 26, 30, 31*). Considering the observed anti-inflammatory effects of Kyn-CKA, it was hypothesized that the upregulation of the kynurenine pathway in SCD constitutes a feedback mechanism to minimize the occurrence of vaso-occlusive events through Kyn-CKA formation. To test this concept, SCD mice were supplemented with Kyn-CKA and the formation of LPS-induced microvascular obstructions in the pulmonary circulation was quantified in real-time by intravital microscopy. Consistent with this hypothesis, increasing systemic Kyn-CKA levels before LPS challenge significantly reduced the size of neutrophil-platelet aggregates and attenuated microvascular occlusions. This effect was likely due to combined inhibitory effects on endothelial adhesion molecule expression, as well as NF-κB and NLRP3-dependent pathway engagement in myeloid cells and platelets.

Importantly, and despite the anti-inflammatory effects of Kyn-CKA, significant associations have been found between increased plasma kynurenine levels and poor outcomes in different human diseases (*6–9*). However, while these findings have often led investigators to assume that kynurenine synthesis upregulation is a pathological process, evidence suggests that these responses are a part of a compensatory anti-inflammatory reaction to the specific pathological condition (*10, 32*). For instance, while genetic IDO1 ablation potentiates antitumor activity and inhibits metastasis (*33*), this same intervention worsens autoimmune disease (*34*). Furthermore, IDO1 gene transfer enhances tissue engraftment and survival in an allogenic transplantation model (*35*), and exogenous kynurenine administration is protective against mortality due to severe endotoxemia (*2*). Considering the present results, we propose that the upregulation of the kynurenine pathway is an adaptive signaling response that suppresses inflammation and protects tissues from injury via Kyn-CKA formation.

In summary, this report provides unequivocal evidence that Kyn-CKA is an endogenous mediator that contributes to the immunoregulatory effects of kynurenine and its metabolites, thus impacting disease progression in chronic inflammatory conditions.

## Supporting information

VOC-Control 1

VOC-Control 2

VOC-Control 3

VOC-Control 4

VOC-Control 5

VOC-Kyn-CKA 1

VOC-Kyn-CKA 2

VOC-Kyn-CKA 3

VOC-Kyn-CKA 4

VOC-Kyn-CKA 5

## Author contributions

Conceptualization: DAV, MC, MFP, GJK, PS, FJS

Methodology: DAV, MC, MFP, SRS, SGW, TB, PS

Investigation: DAV, MC, MFP, SRW, TB, SG, MD, NC, SY

Funding acquisition: DAV, SOF, BAF, SOF, FJS, ACS, PS

Project administration: DAV

Resources: DAV, GJK, SG, SGW, SOF, BAF, FJS, PS

Supervision: DAV, ACS, PS, GJK, SOF, FJS, BAF

Writing – original draft: DAV, MC, MFP

Writing – review & editing: DAV, BAF, PS, FJS

## Data and materials availability

All data are available in the main text or the supplementary materials.

## Supplementary Materials

Materials and Methods

Figs. S1 to S4

Movies Control-1 to 5, and Kyn-CKA-1 to 5

