## Supplementary Data for "Immunomodulatory actions of a kynurenine-derived endogenous electrophile"

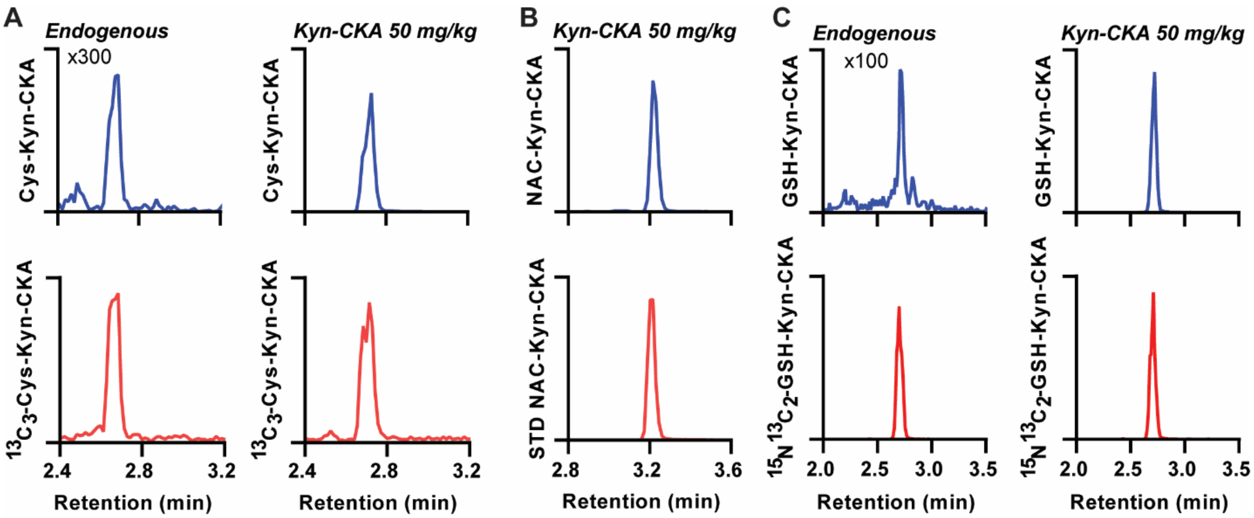


**Supplementary Fig. 1. Kyn-CKA metabolites are present *in vivo*.** (A) Representative LC-MS/MS traces showing urinary Cys-Kyn-CKA (MRM 311/120) in the absence and 8 h following intraperitoneal Kyn-CKA co-eluting with an isotopically labeled standard (MRM 314/123). The x300 factor is relative to the area of the Kyn-CKA supplemented trace. (B) Representative LC-MS/MS trace of urine NAC-Kyn-CKA (MRM 353/162) 8 h post Kyn-CKA co-eluting with a synthetic standard. (C) Representative LC-MS/MS traces of hepatic GSH-Kyn-CKA adducts in the absence and presence of Kyn-CKA supplementation at 8 h. The x300 factor is relative to the area of the Kyn-CKA supplemented trace.

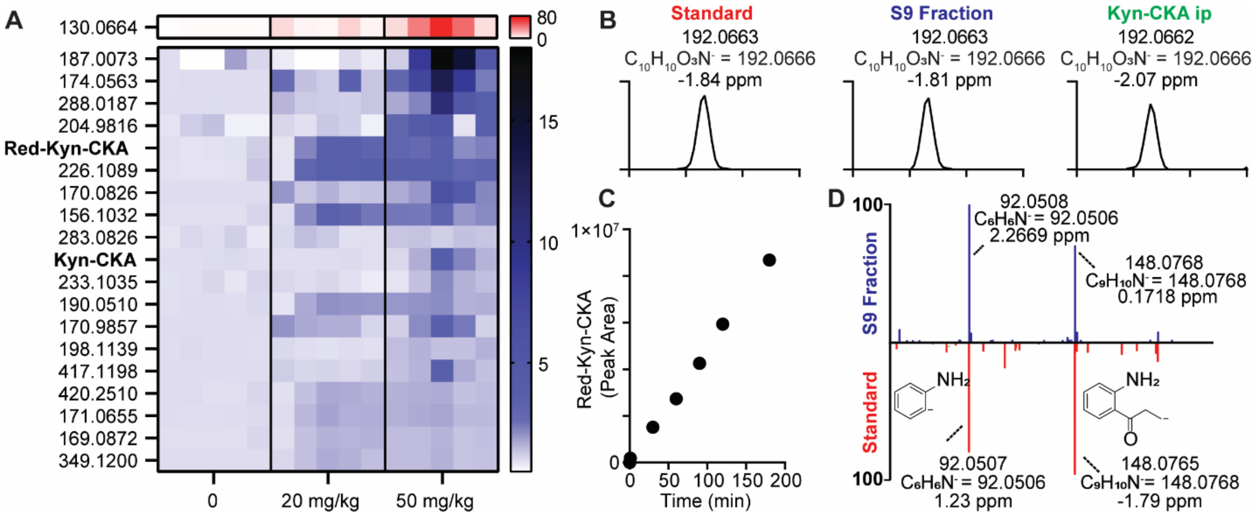


**Supplementary Fig. 2. Candidate metabolites modulated by Kyn-CKA administration.** (A) Heatmap showing the 20 top features detected by untargeted LC-HRMS to be increased in mouse plasma following Kyn-CKA treatment using the free XCMS suite. (B) Confirmation of molecular composition (at the 2 ppm level), using high-resolution mass spectrometry, for synthetic Red-Kyn-CKA, the product of S9 fraction reduction of Kyn-CKA and Red-Kyn-CKA detected in the plasma from Kyn-CKA treated mice at 8 h. (C) Kinetic trace showing the enzymatic conversion of Kyn-CKA (200 μM) to Red-Kyn-CKA by a rat liver S9 fraction (1 mg) in the presence of NADPH (1 mM) at 37°C, pH 7.4. (D) Structural confirmation of S9-derived products by $MS^2$ using synthetic Red-Kyn-CKA.

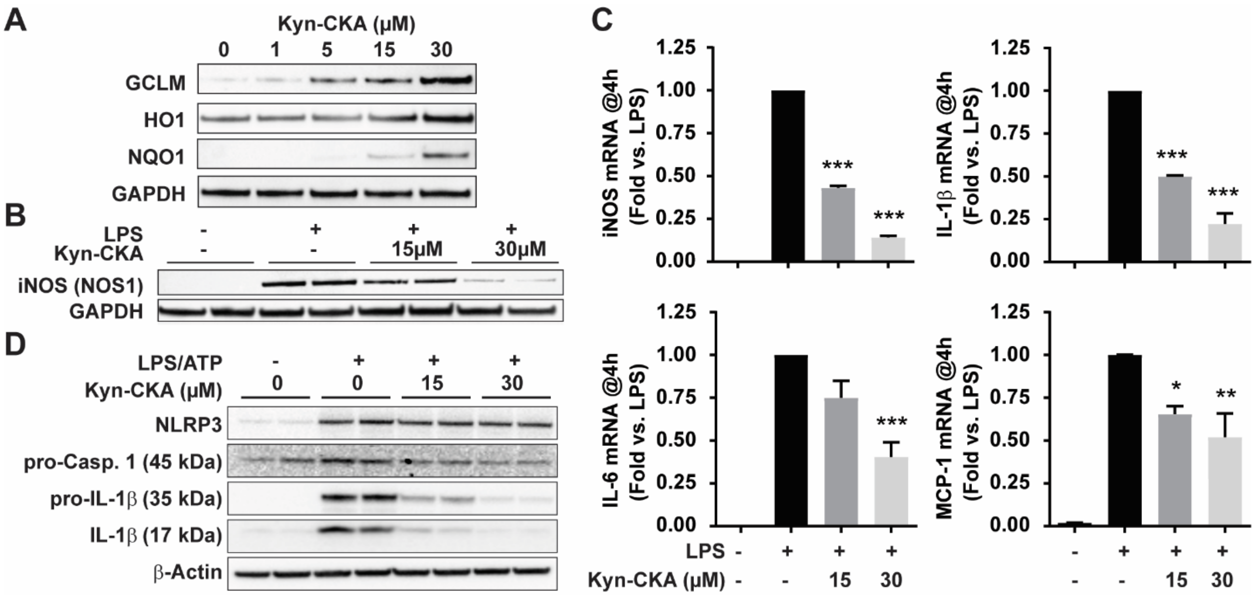


**Supplementary Fig. 3. Kyn-CKA induces Nrf2-dependent proteins and inhibits NF-κB and NLRP3 inflammasome engagement in BMDM** (**A**) Dose-dependent induction of Nrf2 target proteins by Kyn-CKA at 24 h in bone marrow-derived macrophages (B) Kyn-CKA inhibits LPS-induced (100 ng/mL) iNOS expression at 8 h, and (C) iNOS and pro-inflammatory cytokine RNA expression at 4 h, *$p < 0.05$, ** $p < 0.01$, *** $p < 0.0001$ by one-way ANOVA and Tukey's test (n=3). (D) Kyn-CKA inhibits pro-caspase-1 and pro-IL-1β expression, and pro-IL-1β processing in BMDM treated with LPS (100 ng/mL) and ATP (2 mM) for 8 h.

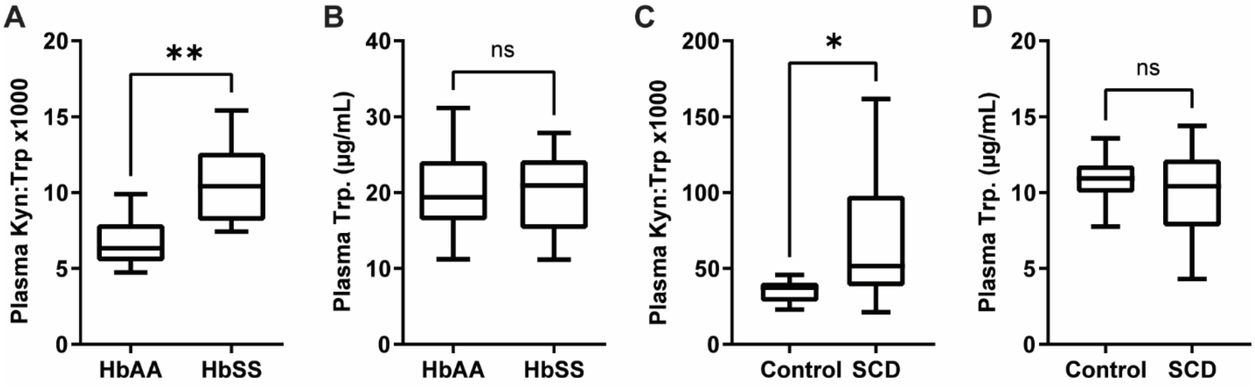


**Supplementary Fig. 4. Kynurenine synthesis is upregulated in murine and human SCD** (A) Plasma kynurenine:tryptophan ratios and (B) tryptophan levels in HbAA (n = 13) and HbSS (n = 8) Townes mice. ** $p < 0.001$ by t-test. (C) Plasma kynurenine:tryptophan ratios and (D) tryptophan levels in control (n = 12) and SCD (n = 10) human donors. * $p < 0.05$ by t-test.
