## Supplementary material for "Immunomodulatory actions of a kynurenine-derived endogenous electrophile": Materials and Methods

**Materials -** *Antibodies:* NLRP3 (Cat# 15101), IL-1β (Cat# 12242), iNOS (Cat# 13120) and GAPDH (Cat# 2118) were purchased from Cell Signaling (Danvers, MA). HO1 (Cat# ADI-SPA-895) was from Enzo Life Sciences (Farmingdale, NY). NQO1 (Cat# ab34173) and VCAM-1 (Cat# ab134047) were from Abcam (Cambridge, UK). GCLM (Cat# 14241-1-AP) was from Proteintech (Rosemont, IL). *Primers:* IL-1β (Mm00434228), IL-6 (Mm00446190), MCP-1 (Mm00441242), HO1 (Mm00516005), NQO1 (Mm01253561), GCLM (Mm01324400) and iNOS (Mm004405021) were from Thermo Fisher (Waltham, MA). Kim-1 (Cat# 12001950) was from BioRad (Hercules, CA) and Actin (Cat# 4351315) from Applied Biosystems (Waltham, MA). *Reagents:* Probenecid was from Enzo Life Sciences, L-Tryptophan-$d_3$ and $^{13}C_3$-Cysteine were from Toronto Research Chemicals (Toronto, Canada), Interferon-γ was from BD Pharmingen (Franklin Lakes, NJ), $^{13}C_2$,$^{15}$N-Glutathione was from Sigma-Aldrich (St. Louis, MO), Kyn-CKA (4-(2-aminophenyl)-4-oxobut-2-enoic acid) was custom-synthesized by Toronto Research Chemicals. $^{15}N^{13}C_2$-Glutathione- and $^{13}C_3$-Cysteine- conjugates of Kyn-CKA were generated by reacting 100 equivalents of thiol with Kyn-CKA in 5 mM ammonium bicarbonate pH 8.0 for 1 h at 37°C followed by solid phase extraction using HyperSep C18 cartridges (Thermo Fisher). L-Kynurenine and N-formyl-kynurenine stocks were treated with 35 mg of 3-mercaptopropyl-functionalized silica from SiliCycle (Quebec City, Canada) for 15 minutes at room temperature to remove potential Kyn-CKA traces followed by 0.22 µm filtration. Organic solvents were LC/MS-grade (Thermo Fisher) and all other chemicals were of analytical grade and obtained from Sigma unless specified.

**Chemical syntheses -** *General Methods:* All glassware was oven-dried before use and reactions were performed under an atmosphere of dry $N_2$. Acro-seal dry solvents stored over molecular sieves were purchased and used as received. Flash chromatography was performed by a RediSep Combiflash on prepacked cartridges of 40 - 63 µm silica gel. $^1$H and $^{13}$C NMR spectra were performed at 600 MHz and 150 MHz respectively at ambient temperature, and spectra are referenced to residual nondeuterated solvent ($CHCl_3$: 7.26 ppm, 77.00 ppm). Purity of products was assessed by HPLC-UV using C18 reversed phase column (2x100 mm, 5 µm, Phenomenex, Torrance, CA) at 220 nm.

*Red-Kyn-CKA [(E)-4-(2-aminophenyl)-4-oxobutanoic acid]:* A culture tube charged with 23 mg (E)-4-(2-nitrophenyl)-4-oxobut-2-enoic acid (0.1 mmol, synthesis described below) and Pd/C (10% w/w, 4 mg) was washed in with 2 mL absolute EtOH. The suspension was stirred and sparged briefly with $N_2$, then a $H_2$ balloon with a 3-way valve was attached via a needle through a septum. The solution was evacuated with house vacuum and refilled with $H_2$ twice and then stirred 1 h under $H_2$ balloon pressure at room temperature. The balloon was refilled when empty (1-2 times). After 1 h the balloon was removed and the suspension filtered through Celite. The solvent was evaporated with a stream of $N_2$, then the residue purified by chromatography (silica gel, dichloromethane+1% HOAc, 0-10% MeOH) and the solvent removed by rotary evaporation to afford 7 mg final product (36%) matching reported spectra (*1*). $^1$H NMR (600 MHz, $CDCl_3$) δ (ppm): 7.75 (d, $J$ = 8.0 Hz, 1H); 7.27 (d, $J$ = 8.4 Hz, 1H); 6.66 (d, $J$ = 7.1 Hz, 1H); 6.65 (d, $J$ = 7.9 Hz, 1H); 3.32 (t, $J$ = 6.3 Hz, 2H); 2.76 (t, $J$ = 6.4 Hz, 2H) $^{13}$C NMR (150 MHz, $CDCl_3$) δ (ppm): 199.7, 178.2, 150.3, 134.5, 130.8, 117.4, 117.4, 115.9, 33.6, 28.2.

*(E)-4-(2-nitrophenyl)-4-oxobut-2-enoic acid:* Prepared according to literature procedures and matched to reported spectra (*1, 2*). Briefly, 2-Nitroacetophenone (165 mg, 1.0 mmol) was

charged to a 10 mL culture tube along with 92 mg glyoxalic acid monohydrate (1.0 mmol, 1 eq.) and 2 mL glacial acetic acid. The tube was covered with a septum with a needle inlet and placed in a preheated sand bath (100-110°C), monitored by TLC, for 24-48 h. After 2 d HPLC indicated consumption of starting material. The acetic acid was evaporated with a stream of $N_2$, then the residue purified by chromatography (silica gel, dichloromethane+2% HOAc isocratic). Product fractions were combined and the solvent removed by rotary evaporation to provide 151 mg (68%) pale purple powder. $^{1}$H NMR (600 MHz, $d_6$-DMSO) δ (ppm): 8.24 (d, $J$ = 8.2 Hz, 1H); 7.93 (t, $J$ = 7.5 Hz, 1H); 7.83 (t, $J$ = 7.8 Hz, 1H); 7.71 (d, $J$ = 7.5 Hz, 1H); 7.21 (d, $J$ = 16.0 Hz, 1H); 6.38 (d, $J$ = 16.0 Hz, 1H) $^{13}$C NMR (150 MHz, $d_6$-DMSO) δ (ppm): 192.2, 166.0, 146.2, 138.3, 134.9, 134.4, 134.1, 132.0, 129.0, 124.7.

**Cell culture -** The murine hepatocyte AML-12 cell line (ATCC CRL-2254) was cultured in DMEM:F12 (1:1) supplemented with 10% Fetal Bovine Serum, ITS (10 µg/mL insulin, 5 µg/mL transferrin, and 6.7 ng/mL selenite) (R&D Systems), 40 ng/mL dexamethasone, 100 U/mL penicillin and 100 µg/mL streptomycin at 37 °C and 5% $CO_2$. J774a.1 macrophages (ATCC TIB-67) were cultured in DMEM containing 10% FBS, 100 U/mL penicillin, and 100 µg/mL streptomycin. Murine bone marrow derived macrophages (BMDM) were isolated from tibia and femur of C57BL/6J mice by flushing bone marrow with DMEM high glucose containing 10% FBS. Cells were differentiated over 7 days in complete media (DMEM High Glucose containing 10% FBS and 10% L929 supernatant, 1X non-essential amino acids, 100 µM HEPES, 10 mM glutamine, 1% penicillin/streptomycin and 50 µM β-mercaptoethanol). L929 cells were cultured in DMEM with 10 % FBS + 100 U/mL penicillin and 100 µg/mL streptomycin. Human pulmonary microvascular endothelial cells (HMVEC-L, Lonza, CC2527) were maintained in VascuLife VEGF-Mv medium (Lifeline, LS-1029).

**Real time PCR -** RNA was isolated from cells or pulverized tissues using TRIzol reagent (Invitrogen, USA) (0.4 mL per $1x10^6$ cells or 1 mL/ 100 mg tissue). RNA was quantified by UV absorbance at 260nm and cDNA was obtained by reverse transcription using iScript cDNA kit (Bio-Rad) following the manufacturer's instructions. RT-PCR was performed on StepOnePlus qPCR system using TaqMan Fast Advanced Master Mix (Thermo Fisher) and the primers listed above. All data were calculated using the relative quantification method (ΔΔCt) and results were expressed as the ratio of the gene of interest to actin expression. *Nrf2-dependent targets*: Cells were seeded in a 12-wells plate in complete media ($0.5x10^6$/well). After reaching confluence, cells were treated with Kyn-CKA (0-60 µM) in complete media without FBS for 8 h at 37°C, 5% $CO_2$ followed by RNA isolation and RT-PCR analysis as described. *NF-κB-dependent targets*: J774a.1 macrophages or BMDM were seeded in 12-wells plates in complete media. Confluent cells were pre-incubated with Kyn-CKA (15-60 µM in complete media without FBS) for 40 min, followed by LPS challenge (E. coli O111:B4, 1 µg/mL) for 4 or 8 h at 37°C, 5% $CO_2$. Cells were collected in TRIzol reagent and processed for RT-PCR as before.

**Protein Immunoblotting -** Cultured cells were washed twice with cold PBS, harvested in 1 mL of PBS, and lysed in 200 µL RIPA buffer containing EDTA-free protease (ThermoFisher) and phosphatase inhibitors (Millipore). Flash-frozen and pulverized tissues were resuspended in RIPA buffer (50 mg in 500 µL) containing protease and phosphatase inhibitors as before. Protein concentrations were determined using a BCA Protein Assay Kit (ThermoFisher) and equal amounts of proteins were subjected to SDS-PAGE using 4–12% sodium dodecyl sulfatepolyacrylamide gels. Proteins were transferred to nitrocellulose membranes, blocked with casein in TBS for 1 h at room temperature, and incubated with primary antibodies overnight at 4 °C followed by incubation with HRP-linked secondary antibodies for 1 h, RT. Blots were developed using Clarity Western ECL Substrate in a ChemiDoc XRS+ imaging system (Bio-Rad). Band intensities were quantified using ImageLab 5.0 (BioRad) and normalized to the housekeeping protein GAPDH unless otherwise specified. *Nrf2-dependent protein expression:* Cells were seeded in 6-wells plates ($1x10^6$ cells/well) in complete media until confluence followed by incubation with Kyn-CKA (0 – 60 µM) in complete media without FBS for 16 h. *iNOS expression and NLRP3 engagement*: J774a.1 or BMDM ($1x10^6$ cells/well) were pre-treated with Kyn-CKA (15-60 µM) for 40 min, followed by LPS addition (1 µg/mL, 8 h). For NLRP3 inflammasome studies, ATP (2 mM) was added in the last 30 minutes of the incubation. *Human pulmonary microvascular cells.* Cells were seeded in a 12-well plate, allowed to grow to confluence and treated with 100 ng/ml LPS in the presence of 0-60 µM Kyn-CKA for 16h.

**Plasma hemin determinations -** Hemoglobin derived species were measured in untreated plasma by nonlinear deconvolution analysis of visible spectra between 520 and 700 nm using Microsoft Excel (*3*). Deconvolution spreadsheets were obtained from Dr. Rakesh P. Patel form the University of Alabama at Birmingham.

**Extraction of kynurenine and Kyn-CKA metabolites -** *Cell-based assays:* Pellets from Kyn-CKA (60 µM in HBSS, 37 ºC) or Tryptophan (0.5, 5 mM, 8 h, 37ºC in HBSS) treated cells were washed and lysed in 0.5 mL MeOH in presence of 100 nM Trp-$d_3$. Lysates were then centrifuged, and supernatants dried under nitrogen, followed by resuspension in 100 µL of 10 % methanol. Media samples were diluted 1:100 in 10% MeOH in the presence of 500 nM Trp-$d_3$. When needed, cells were pre-incubated with probenecid for 1 h to block MRP function. *In vivo assays*: Urine (50-200 µL) and plasma (100 µL) samples were deproteinized by addition of 1 mL cold acetonitrile in presence of Trp-$d_3$ internal standard. Samples were incubated for 10 minutes at -80°C, followed by centrifugation, solvent evaporation and reconstitution in 10% MeOH. For tissues, frozen pulverized samples were resuspended in 250 µL water (100 mg/mL) and extracted in 1.5 mL of cold acetonitrile followed by two cycles of freezing and thawing. Supernatants were dried under nitrogen and reconstituted in 100 µL water. Trp-$d_3$ was used as an internal standard at a final concentration of 500 nM.

**LC-MS/MS analyses -** *High Resolution-MS:* Samples were resolved using a reversed-phase HPLC column (2 x 100 mm 5 µm Luna Phenyl-Hexyl 100 Å column, Phenomenex) and a flow rate of 0.45 mL/min using the following solvent system: A- Water/0.1% Ammonium Acetate/0.2 mM $NH_4F$, B- Acetonitrile/0.01% Formic Acid. Samples were loaded at 3.5% B and eluted with a linear increase to 100% organic solvent over 4.9 minutes. The column was washed with 100% B for 2.3 min and re-equilibrated at 3.5% for an additional 2 min. Samples were analyzed using a Q-Exactive hybrid quadrupole-Orbitrap mass spectrometer equipped with a HESI II electrospray source (Thermo Scientific). Full mass scan analysis ranged from 100 to 600 *m/z* at 35000 resolution. The mass spectrometer analysis was operated in negative mode, with the following parameters: source voltage 3700 V, spray current 4 µA, auxiliary gas flow 15, sheath gas flow rate 25, sweep gas flow rate 8, S-lens RF level 80, capillary temperature 320 °C. Peak areas were integrated at the MS1 level and normalized to Trp-$d_3$ internal standard. MS2 data was used to confirm metabolite identification. *Triple Quadrupole MS:* Metabolites separation was performed using a reversed-phase HPLC column (2 x 100 mm 5 µm C18 Luna (2) column, Phenomenex) at a flow rate of 0.7 mL/min using the following solvent system: A- Water/0.2 mM $NH_4F$, B- Acetonitrile/0.01% Formic Acid. Samples were loaded at 2% B and eluted with a linear gradient from 2-100% of B over 4 min. The column was then washed with 100% solvent B for 3 min and re-equilibrated at 2% for an additional 5 min. Multiple reaction monitoring mass spectrometry analysis was performed using a Qtrap 6500+ (Sciex) in the negative ion mode with the following parameters: curtain gas 50, ion spray voltage -4500 V, temperature 650 ºC, ion source gas(1) 60, ion source gas(2) 55, collision gas -2. The following transitions were used for detection of metabolites and internal standards: kynurenine (207.08/190.0), N-formyl-kynurenine (235.1/190.0), Kyn-CKA (190.0/128.0, 190.0/144.0, 190.0/146.0), GSH-Kyn-CKA (497.0/306.0), Red-Kyn-CKA (192.1/148.1 and 192.1/92.0), Cys-Kyn-CKA (311.1/120.0 and 311.1/190.0), NAC-Kyn-CKA (369.0/192.0),$^{13}C_3$-Cys-Kyn-CKA (314.1/123.0 and 314.1/190.0) $^{15}N^{13}C_2$-GSH-Kyn-CKA (500.1/309.1), Trp (203.1/116.0) and Trp-$d_3$ (206.1/116.0).

**Plasma cytokines -** Plasma cytokine levels in Kyn-CKA and LPS treated mice were analyzed using a MILLIPLEX MAP Mouse High Sensitivity T Cell Premixed Panel (Millipore) by the UPCI Cancer Biomarkers Facility: Luminex Core Laboratory.

**Animal studies -** All animal experiments were performed with the approval of the Institutional Animal Care and Use Committee of the University of Pittsburgh (19054960, 19116001). *Kynurenine and Kyn-CKA treatments:* Male C57BL/6J mice (RRID:IMSR_JAX:000664), aged 9 -15 weeks, were injected intraperitoneally with L-kynurenine in saline (50 mg/kg) or Kyn-CKA in PBS (20-50 mg/kg) for one or two consecutive days. Some mice were challenged with LPS (10 mg/kg ip) 30 minutes following Kyn-CKA or vehicle. Urine samples were collected in metabolic cages, volumes recorded, and creatinine levels measured using a commercial colorimetric kit (Cayman Chemicals). Plasma and tissues were harvested at the time of sacrifice and flash frozen in liquid nitrogen. All samples were stored at -80ºC. *Intravital lung microscopy:* Male and female humanized "Townes" SCD mice (HbSS, homozygous for $Hba^{tm1(HBA)Tow}$ and $Hbb^{tm2(HBG1,HBB^*)Tow}$, 8 - 12 weeks, RRID:IMSR_JAX:013071) (*4*) were injected with PBS or Kyn-CKA (10mg/kg iv) and then challenged with LPS (0.1 µg/kg iv) 30 minutes later. Imaging was performed 3 h following Kyn-CKA or vehicle as previously (*5*). Mice were anesthetized with ketamine (100 mg/kg ip) and xylazine (20 mg/kg) followed by cannula insertion into the right carotid artery and tracheotomy to facilitate mechanical ventilation and anesthesia (95% $O_2$ plus 1-2% isoflurane). A small portion of the previously exposed left lung was immobilized against a coverslip using a vacuum enabled device. Mice were then injected into the carotid artery catheter with ~125 µg/mouse FITC-dextran, 12 µg/mouse AF546-conjugated Ly6G antibody and 7 µg/mouse V450-CD49b antibody to visualize the pulmonary microcirculation and the presence of infiltrating neutrophils and platelets, respectively. Time-series of two-dimensional fluorescent images (450/20 nm, 525/50 nm and 576/26 nm) were collected in a Nikon multiphoton-excitation microscope at ~15 frames per second following excitation at 850 nm using a resonant scanner. Each FOV was 256 µm x 256 µm with an x-y plane resolution of 0.5 µm per pixel. Imaging was performed for 30 minutes with each FOV monitored for 30 seconds. Vaso-occlusions were assessed in at least 20 FOVs per mouse and quantified as the average number of vaso-occlusions per FOV and the average area of each individual vaso-occlusion.

**Human subjects** – Plasma and urine samples from male and female donors were aliquoted and stored at -80$^{o}$C within 1 hour of collection (University of Pittsburgh IRB CR19030018). Additional age- and race-matched samples from healthy volunteers were obtained from Innovative Research (Novi, MI). Control plasma samples (n=12) were 50% female, median age 35 (range: 21-62 years), 100% African descent. SCD plasma samples were 60% female, median age 39 (24-62), 90% African descent, 10% white, 60% HbSS, 2% HbSC, 10% HbSB+, 10% HbSB0. Control urine samples (n=8) were 50% female, median age 40 (range: 27-62 years), 100% African descent. SCD urine samples were 71% female, median age 41 (24-62), 100% African descent, 71% HbSS, 29% HbSC.

**Statistical Analysis -** All statistical analysis was performed using Prism 9.1.2 (GraphPad, San Diego, CA). Data were expressed as mean ± SD (standard deviation, shown as error bars) and the distributions tested for normality. Differences across normal data populations were examined by either Student's t-test or one way-ANOVA followed by Tukey's multiple comparison test unless otherwise specified. Non-parametric tests were utilized for non-normal data distributions. Differences between groups with $p < 0.05$ were deemed significant. Figures show data derived from technical replicates or individual animals and are representative of at least two consistent experiments. Data outliers were identified using the ROUT method with a maximum false discovery rate (Q value) of 1%. No exclusion criteria were pre-established.

**Supplementary Table 1 - Antibodies:**

| Antibody | Catalog number | Company | RRID |
|---|---|---|---|
| NLRP3 | 15101 | Cell Signaling | AB_2722591 |
| Pro-IL-1β | 12242 | Cell Signaling | AB_2715503 |
| IL-1β | 12242 | Cell Signaling | AB_2715503 |
| iNOS | 13120 | Cell Signaling | AB_2687529 |
| VCAM-1 | ab134047 | Abcam | AB_2721053 |
| HO1 | ADI-SPA-895 | Enzo Life Sciences | AB_10618757 |
| NQO1 | ab34173 | Abcam | AB_2251526 |
| GCLM | 14241-1-AP | Proteintech | AB_2107832 |
| GAPDH | 2118 | Cell Signaling | AB_561053 |
| α-Rabbit (secondary) | 7074 | Cell Signaling | AB_2099233 |
| α-Mouse (secondary) | 7076 | Cell Signaling | AB_330924 |

**Supplementary Table 2 - Primers used for RT-PCR:**

| Primer | Catalog number | Company |
|---|---|---|
| IL-1β | Mm00434228 | ThermoFisher |
| IL-6 | Mm00446190 | ThermoFisher |
| MCP1 (Ccl2) | Mm00441242 | ThermoFisher |
| HO1 (Hmox1) | Mm00516005 | ThermoFisher |
| NQO1 | Mm01253561 | ThermoFisher |
| GCLM | Mm01324400 | ThermoFisher |
| iNOS (Nos2) | Mm00440502 | ThermoFisher |
| Kim-1 (Havcr1) | 12001950 | BioRad |
| Actin | 4351315 | Applied Biosystems |

**Supplementary Table 3 - Transitions for MRM analysis:**

| Metabolite | Q1 | Q3 | DP | EP | CE | CXP |
|---|---|---|---|---|---|---|
| Kynurenine | 207.1 | 190.0 | -65 | -8 | -22 | -10 |
| N-Formyl-kynurenine | 235.1 | 190.0 | -65 | -8 | -22 | -10 |
| Kyn-CKA | 190.0 | 128.0 | -65 | -5 | -15 | -8 |
| Kyn-CKA | 190.0 | 144.0 | -65 | -8 | -22 | -12 |
| Kyn-CKA | 190.0 | 146.0 | -65 | -8 | -15 | -10 |
| GSH-Kyn-CKA | 497.0 | 306.0 | -55 | -10 | -20 | -8 |
| Red-Kyn-CKA | 192.1 | 148.1 | -55 | -7 | -15 | -6 |
| Red-Kyn-CKA | 192.1 | 92.0 | -55 | -3 | -27 | -12 |
| Cys-Kyn-CKA | 311.1 | 120.0 | -70 | -5 | -30 | -10 |
| Cys-Kyn-CKA | 311.1 | 190.0 | -70 | -5 | -25 | -10 |
| NAC-Kyn-CKA | 369.0 | 162.0 | -70 | -8 | -20 | -7 |
| $^{15}N^{13}C_2$-GSH-Kyn-CKA | 500.1 | 309.1 | -70 | -5 | -20 | -10 |
| $^{13}C_3$-Cys-Kyn-CKA | 314.1 | 123.0 | -70 | -5 | -30 | -10 |
| $^{13}C_3$-Cys-Kyn-CKA | 314.1 | 190.0 | -70 | -5 | -25 | -10 |
| Trp-$d_3$ | 206.1 | 116.0 | -60 | -8 | -22 | -8 |

DP: declustering potential, EP: entrance potential, CE: collision energy, CXP: collision cell exit potential
